# Four Phases of a Force Transient Emerge from a Binary Mechanical System

**DOI:** 10.1101/2023.09.20.558705

**Authors:** Josh E. Baker

## Abstract

Models of muscle contraction are important for guiding drug discovery, drug validation, and clinical decision-making with the goal of improving human health. Models of muscle contraction are also key to discovering clean energy technologies from one of the most efficient and clean-burning machines on the planet. However, these important goals can only be met through muscle models that are based on science. Most every model and mechanism (e.g., a molecular power stroke) of muscle contraction described in the literature to date is based on a corpuscular mechanic philosophy that has been challenged by science for over two decades. A thermodynamic model and mechanisms (e.g., a molecular switch) of muscle contraction is supported by science but has not yet been tested against experimental data. Here, I show that following a rapid perturbation to the free energy of a thermodynamic muscle system, a transient force response emerges with four phases, each corresponding to a different clearly-defined thermodynamic (not molecular) process. I compare these four phases to those observed in two classic muscle transient experiments. The observed consistency between model and data implies that the simplest possible model of muscle contraction (a binary mechanical system) accurately describes muscle contraction.

## INTRODUCTION

Muscle contains a large macromolecular assembly of proteins arranged around actin and myosin filaments within which myosin motors generate force upon binding to an actin filament (Woledge et al. 1985; Finer et al. 1994). High affinity binding of a myosin motor to an actin filament occurs with an intermediate step in the actin-myosin catalyzed ATP hydrolysis reaction cycle (Lymn and Taylor 1971; Goldman 1987; Cooke 1997; Sweeney et al. 2020) that is gated by the release of inorganic phosphate, P_i_ (MDP to AMD in Fig. 1A). Actin binding induces an energetically favorable switch-like lever arm rotation in a myosin motor that displaces compliant elements external to the motor (Fig 1A) (Huxley 1969, 1974; Rayment et al. 1993; Uyeda et al. 1996; Baker et al. 1998; Sabido-David et al. 1998; Llinas et al. 2013). This motor working step is directly observed in single molecule mechanics experiments where upon binding to actin a single myosin motor displaces an actin filament a distance, *d*, of ∼8 nm within a millisecond of binding to actin (Finer et al. 1994; Molloy et al. 1995; Guilford et al. 1997). A steady state binding rate is established by the ATPase reaction through which myosin motors are irreversibly transferred from AMD to MDP following ADP release, ATP binding and ATP hydrolysis (*v* in Fig. 1A). We have shown that in in vitro assays the myosin working step is reversible (Baker et al. 2002) with rate constants (*f*_*+*_ and *f*_*–*_ in Fig. 1A) that depend on an external force (Stewart et al. 2021). In short, myosin is a binary switch that both generates and responds to forces external to the muscle system (Baker et al. 1999; Baker and Thomas 2000; Baker 2004). The fundamental question remains, what is the relationship between this simple molecular mechanism and muscle contraction?

**Figure 1.**
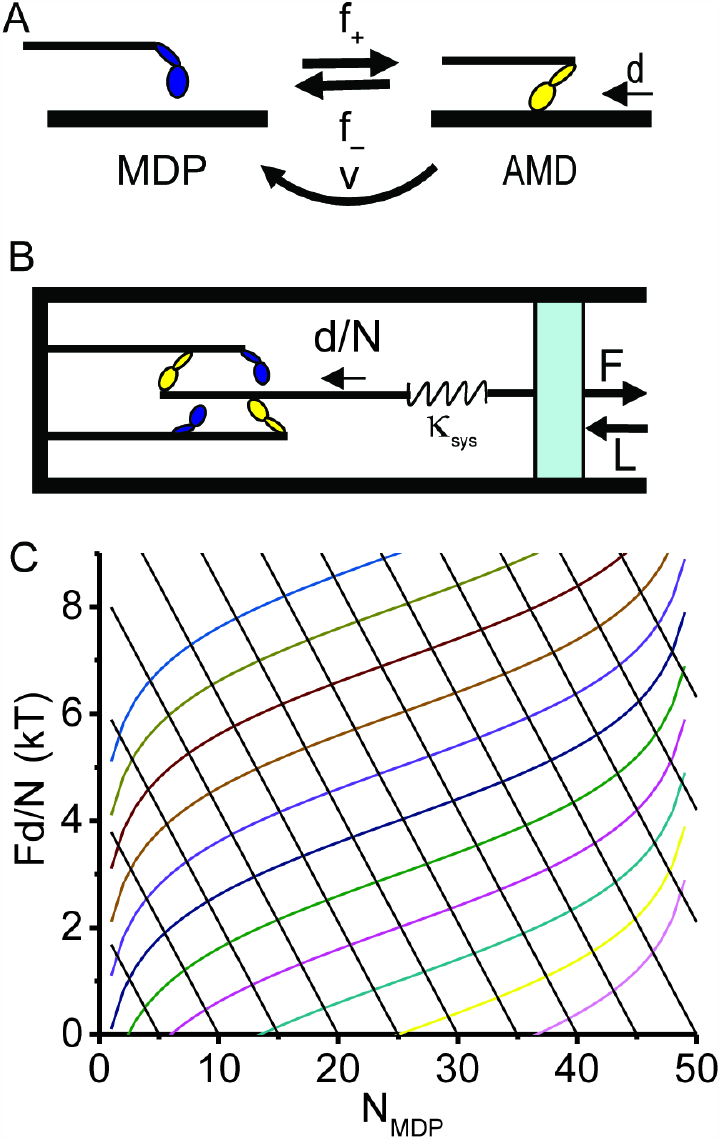
Binary Mechanical Model System. (A) A two state (MDP and AMD) scheme in which a myosin motor, M, with bound ADP, D, and inorganic phosphate, P_i_, undergoes a discrete conformational change upon binding actin, A, and releasing P_i_ to generate a displacement, *d*, at a rate *f*_*+*_. The reverse transition occurs at a rate *f*_*–*_. Through the actin-myosin-catalyzed ATP hydrolysis reaction, AMD is irreverbly transferred to MDP at the ATPase rate, *v*. (B) The laws of mechanics of a binary mechanical system can be described by a single system spring of stiffness *κ*_*sys*_ that on one end (right) describes the macroscopic mechanics (force, *F*, and length, *L*) of muscle and on the other end (left) describes the molecular force, *κ*_*sys*_*d*/*N*, generated with each myosin binding step. (C) Equation 4 (colored curves) is plotted at ΔG° increments of 1 kT, and Eq. 3 is plotted (black lines) at *N*_*MDP*_^*o*^ increments of 5.

In 1938, A.V. Hill proposed a thermodynamic theory of muscle contraction (Hill 1938). In 1957, A.F. Huxley proposed a molecular mechanic theory of muscle contraction (Huxley 1957). The difference between these two theories is fundamental and profound. A.V. Hill’s theory is based on classic chemical thermodynamics, whereas Huxley’s theory, formalized by T.L. Hill in 1974 (Hill 1974), is based on a pre-thermodynamic corpuscular mechanic philosophy where the mechanical properties of the system are determined by the mechanical properties of the molecules within that system.

Despite 200 years of scientific challenges to corpuscular mechanics, 85 years of unrefuted evidence that muscle is a thermodynamic system, and thermodynamic mechanisms (e.g., a molecular switch) that are directly observed in muscle and single molecule mechanic studies, almost every mechanism of muscle contraction described in literature to date is based on a corpuscular mechanic model (e.g., a molecular power stroke). This is a problem, considering that thermodynamic muscle models are supported by science and corpuscular mechanic models are not.

Muscle models are important for drug discovery, drug validation, discovery of disease mechanisms, and clinical decision making, and the physicians, researchers, patients, investors and the public who care about these problems want to know that we are using science to solve them. Moreover, muscle is one of the most efficient, clean-burning machines on the planet, and new clean energy technologies can only be learned from muscle if we use science to understand how these machines work. Thus, it is important that we determine which of these theories is incorrect, and this requires further testing a thermodynamic model against experimental data. To that end, here I compare a thermodynamic model of muscle force transients with experimental data.

Gibbs describes thermodynamics as the laws of mechanics of a system of molecules as measured by an observer, not the laws of mechanics of the molecules within a system as reasoned by corpuscularians. I have recently shown that the laws of mechanics of an ensemble of myosin motor switches can be described by a single entropic system spring (Fig. 1B), where one end of the spring (Fig. 1B, right) is held at a macroscopic force, *F*, or length, *L*, and the other end of the spring (Fig. 1B, left) is stretched by molecular steps of effective size, *d*/(*a·N*), where *N* is the number of myosin motors and *a* is the extent to which molecular force generation equilibrates with the equilibrium free energy isotherm.

A switch-like lever arm rotation in myosin induced by actin binding is the single molecular mechanical mechanism. When muscle is held at a fixed length (Fig. 1B, right), binding steps generate force (Fig. 1B, left; Fig. 1C, black lines). When muscle shortens isothermally (Fig. 1B, right), binding steps perform work with muscle shortening (Fig. 1B, left; Fig. 1C, colored curves). And when active muscle is held at a fixed force (Fig. 1B, right), binding steps (Fig. 1B, left) both generate force and perform work (Baker 2022a). In short, different thermodynamic mechanisms of muscle contraction emerge from a single binding mechanism.

Here, I consider the chemomechanical response to a rapid perturbation, *δE*, in energy of an equilibrium binary mechanical system. Following *δE*, molecular force generation rapidly reaches a binding isotherm. However, if passive elastic elements contribute to muscle force, molecular force generation equilibrates with a pseudo-equilibrium isotherm before equilibrating. The model thus predicts four phases of a force transient. Phase 1 is *δE*; Phase 2 is molecular force generation toward a pseudo-equilibrium isotherm; Phase 3 is chemical relaxation toward the equilibrium isotherm; and Phase 4 is a return to the initial force along the equilibrium isotherm.

Transient force responses to rapid mechanical and chemical perturbations have been widely studied in muscle with the goal of providing insights into fast muscle kinetics and mechanics (Civan and Podolsky 1966; Huxley and Simmons 1971; Ford et al. 1977; Kawai and Halvorson 1991; Dantzig et al. 1992). In the classic experiments of Huxley and Simmons (Huxley and Simmons 1971), an isometric muscle fiber was mechanically perturbed by rapidly lengthening or shortening the fiber by different distances, resulting in a four-phase force transient response. In the classic experiment of Dantzig et al. (Dantzig et al. 1992), a skinned isometric muscle fiber was chemically perturbed by rapidly changing the phosphate concentration in muscle, resulting in a three-phase force transient response. Using a minimal binary mechanical model, I simulate muscle force transients and compare the simulations to force transients observed in response to both mechanical (Huxley and Simmons) and chemical (Dantzig et al.) perturbations. I show that in all cases the four phases observed are accounted for by the thermodynamic processes described above with simulated rates and amplitudes that are consistent with measured values.

In a binary mechanical model, the laws of mechanics of muscle are described by two simple thermodynamic relationships (a Gibbs free energy equation and a force equation for a system spring). From these physical laws, a minimal number of macroscopic parameters must be determined before molecular mechanisms are inferred. This top-down approach to modeling muscle contraction is fundamentally different from a bottom-up molecular mechanics approach in which macroscopic parameters are determined from the laws of mechanics of molecular mechanical devices conceived by corpuscularians. I have shown that the laws of mechanics of a binary mechanical model accurately describe steady state, force generating, and work loop mechanics of muscle (Baker 2022a). Here I show that these same laws account for the four phases of a muscle force transient as thermodynamic (not molecular) processes. A chemical thermodynamic model opens a new frontier in muscle physiology replacing non-physical rational molecular mechanisms with physical thermodynamic mechanisms that advance, constrain, and dramatically simplify our understanding of systems-level physiology.

## RESULTS

### A Two State Kinetic Scheme

Figure 1A illustrates a simple two-state kinetic scheme in which a myosin (M) motor with bound ADP (D) and inorganic phosphate (P_i_) undergoes a switch-like lever arm rotation induced by actin (A) binding and gated by P_i_ release (MDP to AMD). This binding-induced molecular switch (a working step) reversibly displaces a compliant element external to the motor a distance, *d*, with forward, *f*_*+*_, and reverse, *f*_*–*_, rates (Baker et al. 2002; Stewart et al. 2021; Baker 2022a).

The maximum force generated by muscle is observed to increase linearly with the number, *N*, of myosin motors interacting with an actin filament, which means that myosin motors generate force in parallel. For an equilibrium mixture of *N* parallel force generators, the displacement of the effective system spring by a single working step is *d*/*N* (displacing a single bed spring a distance, *d*, displaces the system of *N* parallel springs a distance *d*/*N*). At equilibrium, the system force, *F*, is distributed among all *N* myosin motors (Baker and Thomas 2000) because all motors (bound and detached) are inextricably part of and equilibrated with the macromolecular assembly to which force is applied. However, in a non-equilibrium (non-ergodic) system, only a fraction, *a*, of motors are equilibrated with the system force, in which case the step size is *d/*(*a*·*N*), where the equilibration factor, *a*, ranges from 1 at equilibrium to 1/*N* when on average only 1 of *N* motors is equilibrated with the system force.

### Binding Free Energy

According to Gibbs, a thermodynamic model of muscle describes the laws of mechanics of muscle as they appear to us. In other words, thermodynamics describes the relationship between the mechanics, energetics, and chemistry of the muscle system (Fig. 1A). These relationships are described by the Gibbs reaction free energy for actin-myosin binding (Fig. 1A) defined at a constant muscle force, *F* (Baker et al. 1999; Baker and Thomas 2000)

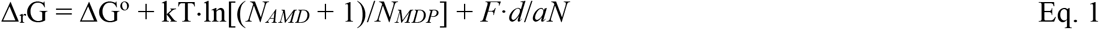

where ΔG° is the standard free energy for the binding reaction in Fig. 1A; *N*_*AMD*_ and *N*_*MDP*_ are the number of myosin motors in the AMD and MDP states (the total number of myosin motors is *N* = *N*_*AMD*_ + *N*_*MDP*_); and the logarithmic term is the change in system entropy with a chemical step from {*N*_*MDP*_, *N*_*AMD*_} to {*N*_*MDP*_–1,*N*_*AMD*_+1} along the system reaction coordinate (Fig. 1C, y-axis, right to left) (Baker 2022a). Here, I approximate (*N*_*AMD*_ + 1) as *N*_*AMD*_ and assume fixed concentrations of actin and basal inorganic phosphate, P_i_, are implicit in ΔG°.

### | A Muscle Spring

The laws of mechanics of a muscle system can be described in terms of a single linear system spring (Fig. 1B) of stiffness, *κ*_*sys*_ (Baker 2022a). One end of the spring (Fig. 1B, right) defines the macroscopic mechanical state of muscle (force, *F*, and length, *L*), while the other end of the spring (Fig. 1B, left) is stretched by molecular binding steps that generate force through *κ*_*sys*_· *d*/*aN* increments. In isometric muscle held at a fixed length (the right side of the spring in Fig. 1B is fixed), no energy is lost to the surroundings as shortening heat or work, which is to say that molecular force generation is adiabatic. In this case force, *F*, increases linearly with the number of bound myosin motors, *N*_*AMD*_, or decreases linearly with the number of detached myosin motors, *N*_*MDP*_, as

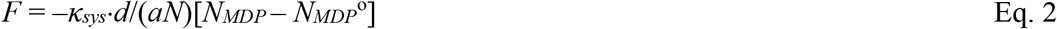

where *N*_*MDP*_^*o*^ is *N*_*MDP*_ at *F* = 0. Multiplying both sides of Eq. 2 by *d/N*

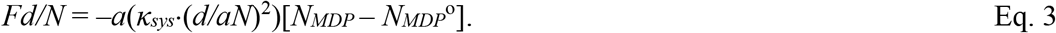

Equation 3 is plotted as *Fd/N* versus *N*_*MDP*_ in Fig. 1C (solid black lines with *N*_*MDP*_^*o*^ increments of 5).

Adiabatic (isometric) molecular force generation (Eq. 2) in the system spring continues until the force, *F*, reaches a binding isotherm (Eq. 1), which occurs when

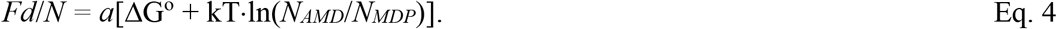

Equation 4 is plotted as *Fd/N* versus *N*_*MDP*_ in Fig. 1C (colored curves with ΔG° increments of 1 kT).

Equations 3 and 4 are two different definitions of system force, *F*, distinguished by whether changes in muscle force occur adiabatically (Fig. 1C, left side) or isothermally (Fig. 1C, right side). In an ideal system (adiabatic or isothermal) the binding reaction follows one or the other of these pathways (black lines or colored curves in Fig. 1C). However in general these two processes do not occur independent one another because under non-adiabatic, non-equilibrium conditions, the force of the system spring must be the same on both sides of the spring (*F* must be the same in both Eqs. 3 and 4). Formally, this requirement is satisfied because the kinetics of molecular force generation (Eq. 3) are defined by energetics (Eq. 4) as

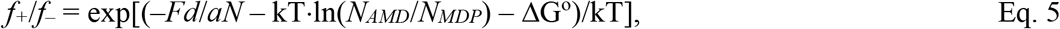

where Eq. 5 is simply the right-hand side of Eq. 1 rewritten to describe the probability of the muscle system being found in state {*N*_*MDP*_,*N*_*AMD*_} relative to {*N*_*MDP*_–1,*N*_*AMD*_+1} along a system reaction energy landscape that is tilted by the energy terms in Eq. 5 (Baker 2022a).

When *a* = 1, molecular force generation stalls along the equilibrium binding isotherm (Δ_r_G = 0) at a force, *F*_*o*_ = –NΔG°/*d* – NkT·ln(*N*_*AMD*_/*N*_*MDP*_). When *a* < 1, molecular force generation stalls along a pseudo-equilibrium (Δ_r_G < 0) binding isotherm at a force, *F* = *a*·*F*_*o*_. There are many mechanisms by which *F* stalls along a pseudo-equilibrium isotherm. During isotonic contractions, muscle is held at a fixed force, *F*, and molecular force generation stalls at a pseudo-equilibrium isotherm, *F* = *a*·*F*_*o*_, where the remaining unequilibrated binding free energy Δ_r_G = (1 – *a*)NΔG° /*d* is lost from the shortening muscle system as heat and work. This is the energetic basis for A.V. Hill’s muscle force-velocity relationship (Hill 1938). Similarly, in partially activated muscle, *a* is the fractional activation. In both examples, a*F*_*o*_ is an internal force balanced against the external force, *F*. During unloaded muscle shortening, internal forces generated between myosin motors prevent equilibration with an external force, *F*, of zero, and values for *a* = 0.2 to 0.3 measured in muscle (Woledge et al. 1985) describe the magnitude of internal forces generated relative to *F*_*o*_ (Baker 2022a). In this example, *F* < *aF*_*o*_. Below, I consider a case in which passive compliant elements contribute to a non-equilibrium internal force, *aF*_*o*_, that transiently exceeds *F*_*o*_, or *F* > *aF*_*o*_.

If the internal energy of a system is perturbed by *δE*, the system energy landscape is further tilted by *δE* along the system reaction coordinate, perturbing the kinetics of force generation from Eq. 5 as

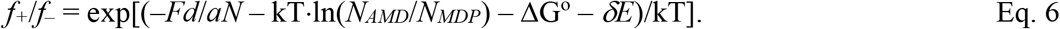

For each energy term, *E*, in Eq. 6 the *E*-dependence of exp(*E*/kT) can be partitioned between forward, *f*_*+*_(*E*), and reverse, *f*_*–*_(*E*), rate constants through a coefficient, *α*_*E*_, that describes the fractional change in *E* prior to the activation energy barrier (Hille 1987). For example, when *δE* and *F* are zero,

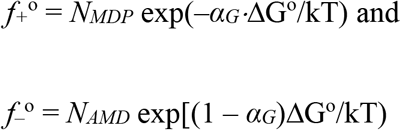

are unloaded rates.

The time courses for all processes, both ideal (Eqs. 3 and 4) and non-ideal (intermediate between Eqs. 3 and 4), are described by three simple master equations. The rate of the two-state reaction in Fig. 1A is

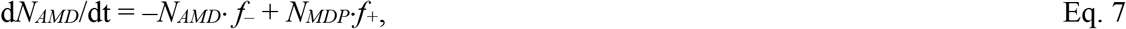

which according to Eq. 2 generates force in the system spring at a rate

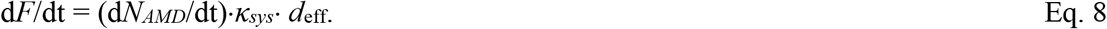

If force generation through Eq. 8 stalls at a pseudo-equilibrium isotherm (i.e., *a* is not equal to 1), the system subsequently equilibrates at a rate *b* (i.e., the rate at which *a* approaches 1), defining the third master equation

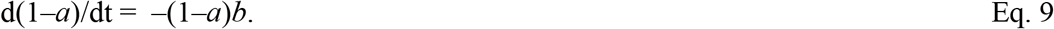

For each mechanism by which equilibration of force generation is prevented (e.g., *a* < 1), there is a unique rate, *b*, at which that mechanism relaxes to allow force to equilibrate with the equilibrium isotherm. When *b* = 0, muscle is stuck in a state (e.g., smooth muscle latch) that is the binary mechanical system equivalent of a frustrated spin state (Baker et al. 2003).

From these equations (Eqs. 7 – 9), four phases of a force transient are evident. The initial perturbation, *δE*, from equilibrium is phase 1. Force generation that reaches a pseudo-equilibrium isotherm (Eq. 8) is phase 2. Chemical relaxation toward the equilibrium isotherm (Eq. 9) is phase 3. And a return to the original stall force along the equilibrium isotherm is phase 4. All simulations below are based on these simple master equations (Eqs. 7 – 9) with rate constants defined from first principles (Eq. 6) and parameters defined in Table 1.

### Simulations of Ideal Adiabatic Force Generation and Isotherm Stretch and Shortening

Figure 2 shows simulations of ideal adiabatic and isothermal processes. Adiabatic (fixed length) molecular force generation (phase 2) is simulated in Fig. 2A following a rapid increase (red line) or decrease (blue line) in the the force, Δ*F*, or length, Δ*L*, of the system spring (*δE* = Δ*F*·*d*_*eff*_ = *k*_*sys*_·Δ*L*·*d*_*eff*_) starting from the equilibrium value, *F*_*o*_ = –*N*ΔG°/*d* (phase 1). Slow isothermal lengthening (Fig. 2B, red line) and isothermal shortening (Fig. 2B, blue line) of the system spring are simulated starting from the parameters at the end of phase 2 in Fig. 2A. The simulations in Figs. 2A and 2B (red and blue lines) are replotted in Fig. 2C (solid red and blue lines) as *F*·*d*_*eff*_ versus *N*_*MDP*_ and are overlaid with Eqs. 3 and 4, showing that simulations based on master equations 7 and 8 are consistent with the idealized thermodynamic relationships in Eqs. 3 and 4.

**Figure 2.**
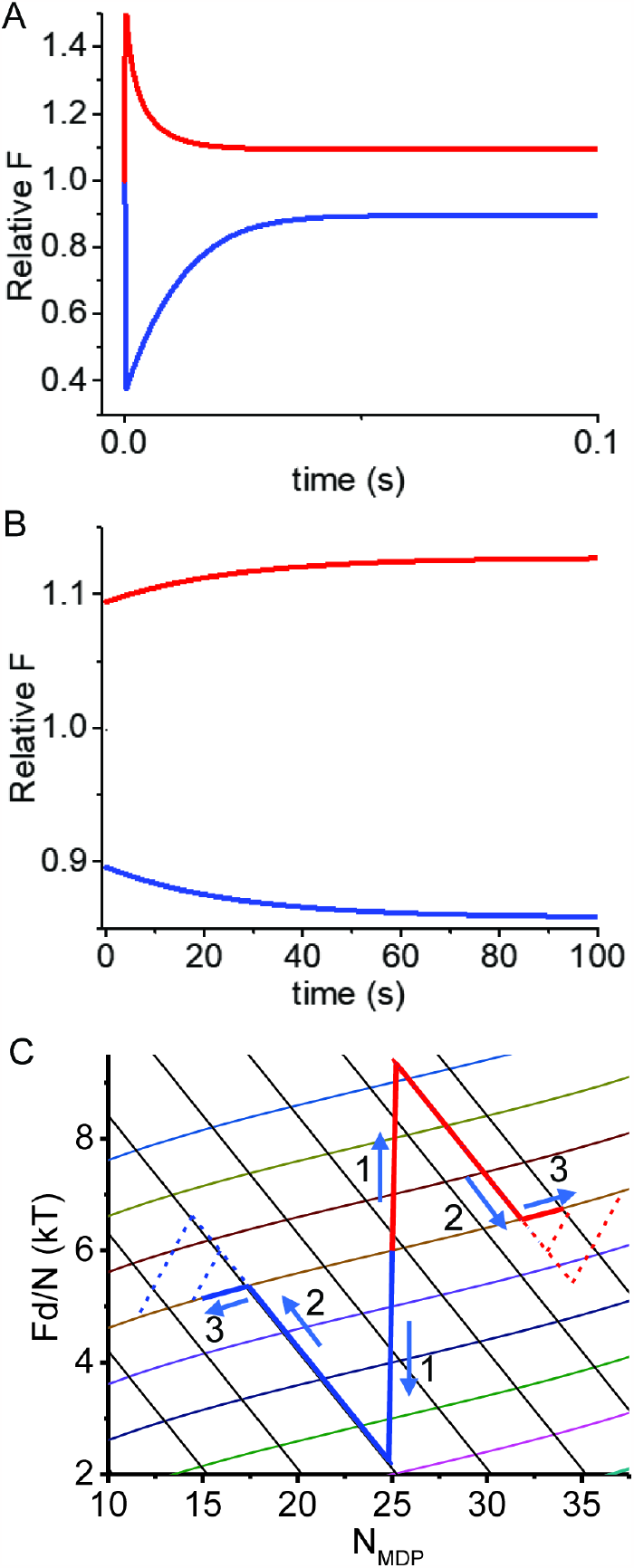
Simulated force transients following rapid length steps, ΔL, using Eqs. 7 and 8 and rate constants defined by Eq. 6. Parameters were initialized to values in Table 1, with *N*_*AMD*_ = *N*_*MDP*_ = 25 and *F*_*i*_ = 41.3 pN (Relative *F*_*i*_ = 1). (A) At t = 0, *F*_*i*_/κ_sys_ was both decreased by Δ*L* = 5.26 nm from a relative *F*_*i*_ of 1 to 0.36 (blue curve) and increased by Δ*L* = 4.74 nm from a relative *F*_*i*_ of 1 to 1.6 (red curve). Force transients initialized at each non-equilibrium *F* value were then simulated. (B) Slow changes in the length of the system spring initialized with equilibrated values for *F, N*_*AMD*_, *N*_*MDP*_ in panel A were simulated. An increase in length was simulated for equilibrated values following the length increase in panel A (red line), and a decrease in length was simulated for equilibrated values following the length decrease in panel A (blue line). (C) Values for *F* and *N*_*MDP*_ from simulations in panels A and B are replotted as *Fd*/*N* vs. *N*_*MDP*_ and overlaid with Eqs. 4 (colored curves) and 3 (solid black lines). The simulations in panels A and B were repeated using values for *z* of 0.4 (low amplitude) and 0.8 (high amplitude) and replotted in panel C (dashed lines).

Figure 2C shows that *F* returns to the initial binding isotherm, which is to say the change in system energy, *δE*, is fully recovered (no energy is lost from the system) through adiabatic force generation. However if the lengthening of the system spring during phase 2 force generation stretches passive parallel elastic elements (e.g., frictional forces), a fraction, *z*, of *δE* is used to perform work on these elements, and *F* approaches a pseudo-equilibrium isotherm that is *z*·*δE* less than the equilibrium isotherm, or *F* = (*F*_*o*_ – *z*·*δE*/*d*) = *aF*_*o*_ and *a* = 1 – (*z*·*δE*/*d*)/*F*_*o*_. Similarly, if phase 2 force generation occurs with the relaxation of strained passive elastic elements, work, *zδE*, is performed by these elements on the system, and *F* approaches a pseudo-equilibrium isotherm that is *z*·*δE* greater than the equilibrium isotherm, where *F* = (*F*_*o*_ + *z*·*δE*/*d*) = *aF*_*o*_ and *a* = 1 + (*z*·*δE*/*d*)/*F*_*o*_. Upon chemical equilibration *F* approaches the equilibrium isotherm (Δ_r_G approaches zero and *a* approaches 1) at a rate *b*.

### Simulations of Adiabatic Force Generation and Mechanical Equilibration

The dotted lines in Fig. 2C are simulations of adiabatic force generation (phase 2) following the same perturbation, *δE*, in Figs. 2A and 2B only here a passive elastic element performs work on the system during adiabatic force generation (values for *z* of 0.4 and 0.8 with a slow mechanical relaxation rate, *b*, of 0.005 s^−1^). Simulations in Fig. 2C (dashed lines) show that phase 2 amplitudes, *F* = *aF*_*o*_, increase with *a* (solid lines are *a* = 1). A return to the equilibrium isotherm (Fig. 2C, dashed lines) resembles isothermal shortening, only here the binding reaction follows the isotherm with a slow loss of *z*·*δE* from the system (phase 3) rather than slow isothermal shortening (muscle length does not change in these simulations).

In Fig. 2C, phases 2 and 3 for all simulations are temporally separate because phase 3 occurs at a relatively slow rate. However, when phase 3 occurs on a time scale comparable to the that of phase 2 (Eq. 3), the two phases begin to merge. Figure 3A is the same simulation shown in Fig. 2C (*z* = 0.4) only here with rates, *b*, of 0, 4, 20, and 40 sec^−1^. These simulations are replotted in Fig. 3B as *F*·*d*_*eff*_ vs. *N*_*MDP*_. When *b* = 0, phase 2 follows Eq. 3 with no phase 3. With an increase in *b*, phase 2 deviates from Eq. 3 until when *b* equals the chemical relaxation rate ((*f*+° + *f*–° = 40 sec^−1^, Table 1), force recovery occurs with a single non-ideal phase intermediate that of adiabatic force generation (Eq. 3) and the isotherm (Eq. 4).

**Figure 3.**
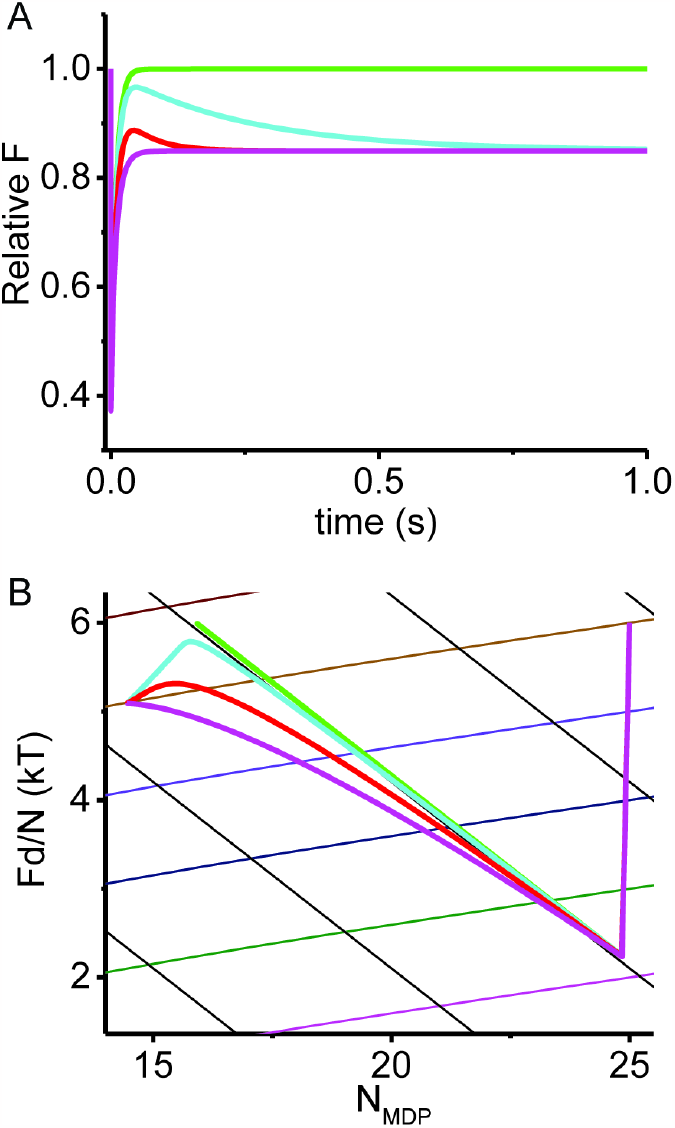
Simulated effects of mechanical equilibration rate, *b*, on force transients. (A) The same simulations performed in Fig. 2 following a rapid shortening step are repeated at *z* = 0.4 and different chemical relaxation rates, *b*, of 0 sec^−1^ (green), 4 sec^−1^ (cyan), 20 sec^−1^ (red), and 40 sec^−1^ (magenta). (B) Values for *F* and *N*_*MDP*_ simulated in panel A are replotted as *Fd*_*eff*_ vs. *N*_*MDP*_ and overlaid with Eqs. 3 (solid black lines) and 4 (colored curves).

The thermodynamic approach to modeling muscle contraction is top-down. That is, a thermodynamic model consists of a limited number of macroscopic parameters (Table 1) that are fit to macroscopic measurements to satisfy the laws of mechanics of the system (Eqs. 3 and 4). Molecular details are then inferred from these fitted thermodynamic parameters. This is a familiar approach when inferring molecular mechanisms from measured changes in ΔG° associated with protein mutations, post-translational modifications, small molecules, and changes in T, pH, and ionic strength. Similarly, the thermodynamic parameter *a* must be determined before molecular mechanisms are inferred from it.

In muscle, the amplitude of phase 2 force generation following a rapid shortening step often overshoots isometric force, demonstrating experimentally that *a >* 1. As described above, one possible mechanism is that strained elastic elements shorten to perform work on the system during phase 2 force generation. In muscle, titin is a large, passive elastic element (Kellermayer et al. 1997) that contributes to muscle force, and any work performed by small decreases in the length of titin during phase 2 force generation contributes to the free energy available for force generation. According to this model, the phase 2 amplitude increases with increasing titin stiffness.

Experimentally, a rapid change in the length of steady state isometric muscle has been shown to elicit a multi-phasic force response. This was first observed in whole muscle (Gasser and Hill 1924; Jewell and Wilkie 1958) and subsequently observed with improved time resolution in single muscle fibers (Huxley and Simmons 1971; Ford et al. 1977). In 1974, Huxley and Simmons observed that when the length of an isometric muscle fiber was rapidly shortened or lengthened the force response occurs in four phases, resembling the four thermodynamic processes described above. Here, I compare computer simulations of these chemical thermodynamic processes with force transients observed by Huxley and Simmons.

Figure 4A shows simulated transient responses to length steps, ΔL, of different amplitudes, which are replotted as *F* vs. *N*_*MDP*_ in Fig. 4B. The transient responses are non-exponential, and so phase 2 rates were determined from 1/t_½_ of the simulated phase 2 where t_½_ is the time at which the force reaches ½ the sum of the maximum phase 1 and phase 3 force. Figure 4C is a plot of simulated phase 2 rates for different length steps overlaid with experimental data obtained from R temporaria muscle at 2° C (Huxley and Simmons 1971).

**Figure 4.**
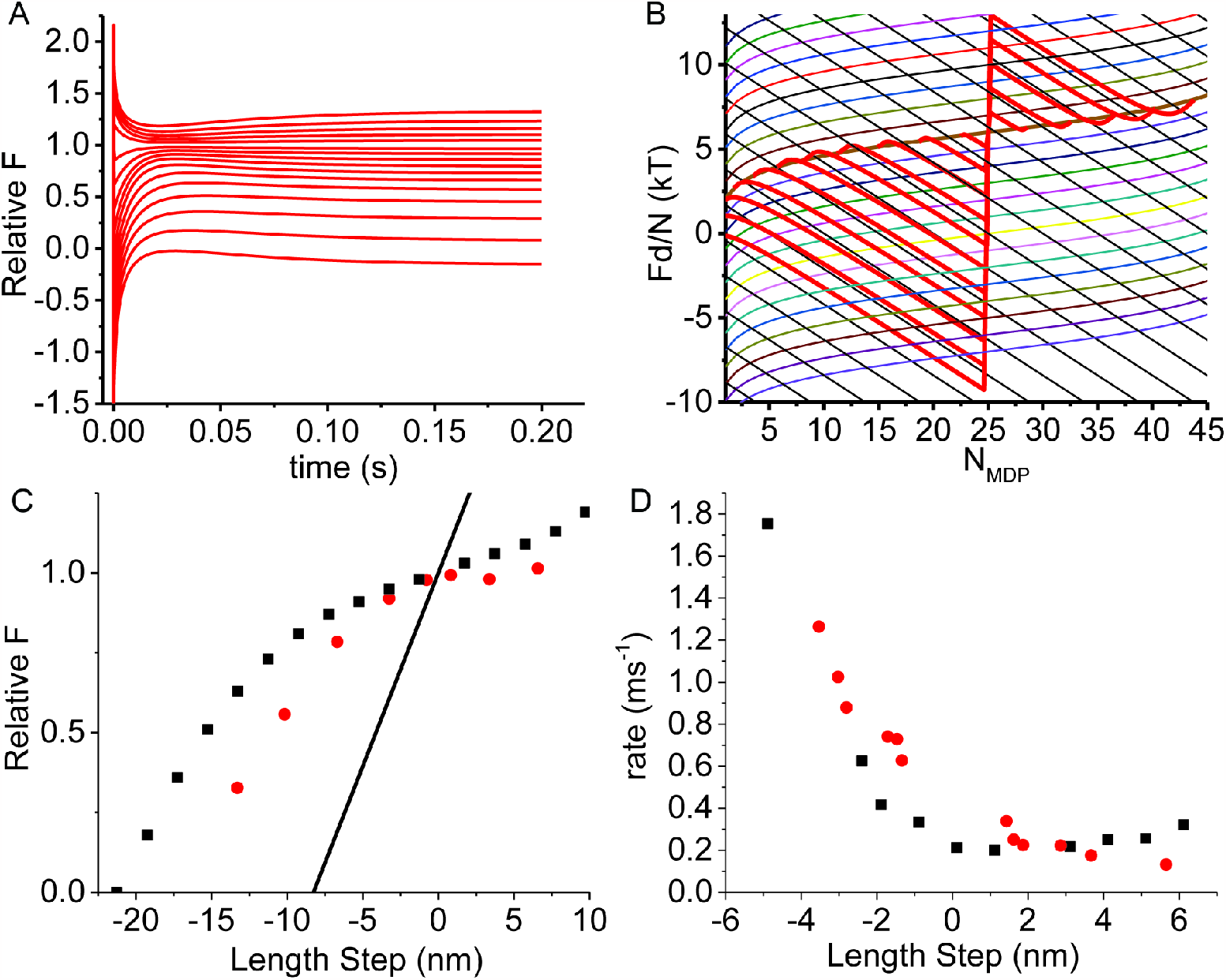
Simulated force transients following length steps, ΔL, using Eqs. 7 and 8 and rate constants defined by Eq. 6. Parameters were initialized to values in Table 1, with *N*_*AMD*_ = *N*_*MDP*_ = 25, *z* = 0.4, *b* = 20 sec^−1^ and *F*_*i*_ = 41.3 pN (Relative *F*_*i*_ = 1). (A) At t = 0, *F*_*i*_/κ_sys_ was increased or decreased by different Δ*L* values followed by simulation as in Fig. 2A (red lines). (B) Values for *F* and *N*_*M*_ simulated in panel A are replotted as Fd/*N* vs. *N*_*M*_ (red lines) and overlaid with Eqs. 3 (solid black lines) and 4 (colored curves). (C) Phase 2 amplitudes were determined from simulations in panel B and plotted (black squares) for different length steps, ΔL, and overlayed with experimental data (red circles) digitized from Huxley and Simmons (Huxley and Simmons 1971). The force, *F*, immediately following the length step, ΔL, (Phase 1 amplitude) is *F* = *F*_*i*_ – ΔL·*k*_*sys*_ and is plotted (black line). (D) Phase 2 rates were determined from simulated transients in panel B as 1/t_½_, plotted (black squares) for different ΔL, and overlayed with experimental data (red circles) digitized from Huxley and Simmons (Huxley and Simmons 1971).

Figure 4D is a plot of the simulated phase 2 amplitudes for different length steps, ΔL, overlaid with experimental data obtained from R temporaria muscle at 2° C (Huxley and Simmons 1971). The mechanism underlying the ΔL dependence of the phase 2 amplitude is geometrically evident from Fig. 4B. Specifically, larger shortening steps, ΔL, require larger increases in *N*_*AMD*_ to equilibrate with the ΔG° isotherm, which decreases the phase 2 amplitude by kTln(N_AMD_/N_MDP_) along the isotherm until at sufficiently large ΔL, *N*_*AMD*_ approaches *N*, and *F* falls off the isotherm, reaching *F* = 0 when *N*_*MDP*_ ° = 0 (Eq. 4).

Force transients have also been measured in muscle following rapid chemical perturbations. For example, Dantzig et al. (Dantzig et al. 1992) showed in skinned muscle fibers that a rapid increase in [P_i_] upon photo-release of caged-P_i_ results in a multi-phasic force response. In these experiments, different final phosphate concentrations, [P_i_]_f_, were achieved through a combination of varying both initial concentrations, [P_i_]_i_, and the amount of caged-P_i_ photo-released. Using an analysis based on the molecular mechanic formalism, they conclude that the initial force response is different from the phase 2 response in a length step experiment and requires even more states (Kawai and Halvorson 1991; Dantzig et al. 1992). Here, I compare the four phases of a mechanical transient in a binary mechanical system with the mechanical response to [P_i_] jumps observed by Dantzig et al.

The myosin working step is associated with the release of P_i_ (Cooke and Pate 1985; Baker et al. 1999, 2002). Thus, a rapid increase in [P_i_] from an initial concentration, [P_i_]_i_, to a final concentration, [P_i_]_f_, increases the system free energy by δE = kT·ln([P_i_]_f_/[P_i_]_i_). Starting from the same initial equilibrium conditions (see Fig. 5 legend) used for the length-step experiments above (only here *f_*° = 0.01 s^−1^) phase 1 is simulated simply by setting δE to kT·ln([P_i_]_f_/[P_i_]_i_). Unlike in length step simulations (Fig. 5), force does not change with this transition, and so phase 1 is not mechanically observed.

**Figure 5.**
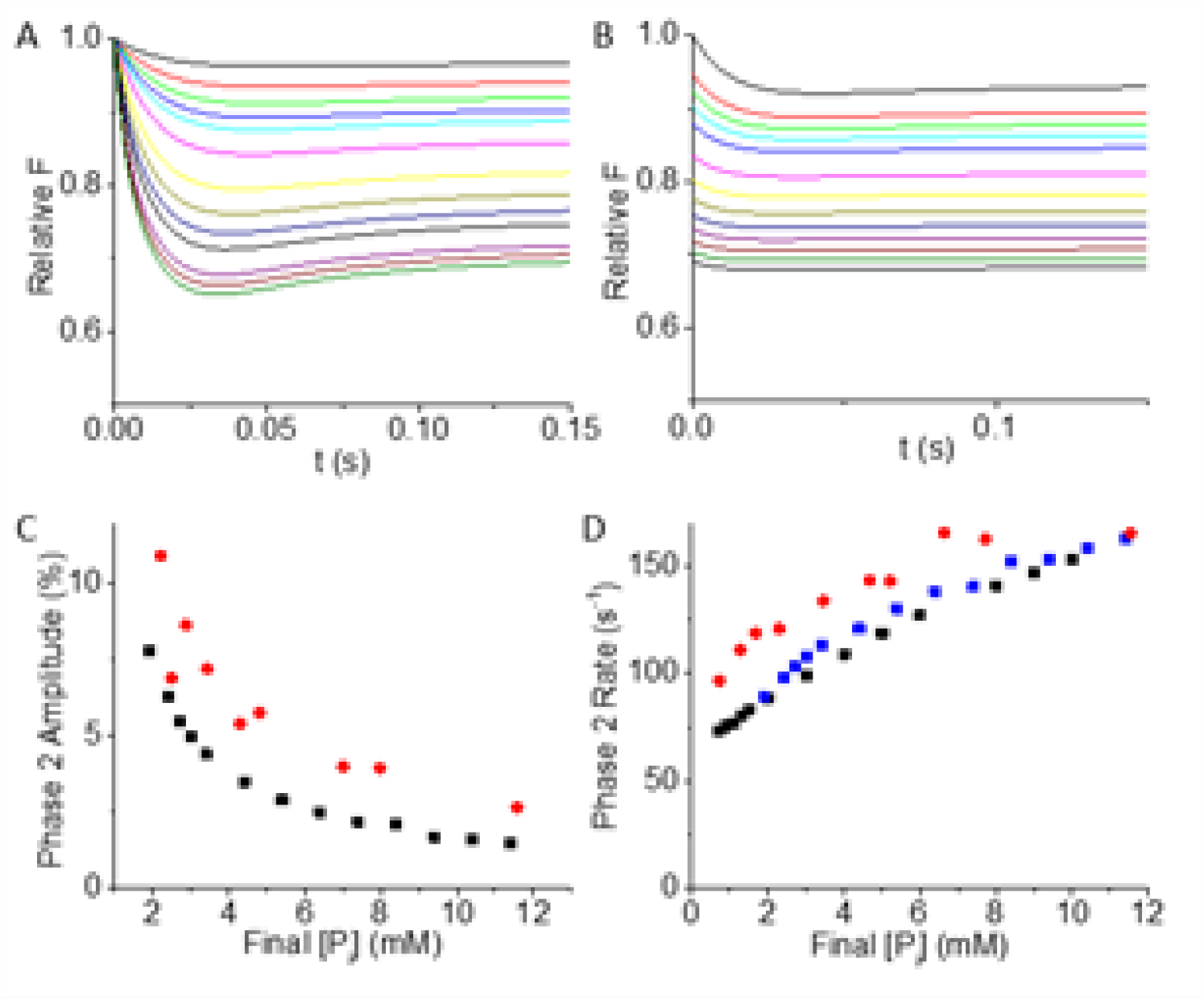
Simulated force transients following rapid increases in [P_i_], using Eqs. 7 and 8 and rate constants defined by Eq. 6. Parameters were initialized to values in Table 1 (except *f_*° = 0.01 s^-1^), *N*_*AMD*_ = *N*_*MDP*_ = 25, *F*_*i*_ = 137.6 pN (Relative *F*_*i*_ = 1), *z* = 0.4, *b* = 20 sec^−1^, and *a*_*dE*_ = 0 ([P_i_] only affects the reverse rate). (A) At *t* = 0, δE = kT·ln([P_i_]_f_/[P_i_]_i_) (Eq. 7), where [P_i_]_f_ and [P_i_]_i_ = 0.5 mM are the final and initial phosphate concentrations. [P_i_]_f_ was set to values ranging from 0.7 to 10 mM (curves top to bottom) and for each [P_i_]_f_ simulations were run (Eqs. 5 and 6) as in Fig. 2A. (B) [P_i_]_i_ was set to values ranging from 0.5 to 10 mM (curves top to bottom) and [P_i_]_f_ = [P_i_]_i_ + 1.4 mM P_i_ and simulations were run as in Fig. 2A. (C) The percent decrease in the phase 2 amplitude relative to the initial force was determined and plotted for different [P_i_]_f_ (black squares) and overlayed with corresponding experimental data (red circles) digitized from Dantzig et al. (Dantzig et al. 1992). (D) The phase 2 rate determined from single exponential fits of phase 2 in both panels A (blue squares) and B (black squares) is plotted and overlayed with experimental data (red circles) digitized from Dantzig et al. (Dantzig et al. 1991).

Figures 5A and 5B show simulated chemical relaxations for different combinations of [P_i_]_f_ and [P_i_]_i_. Because *δE* in Eq. 6 does not discriminate between whether the energy change is chemical or mechanical, the same phases (phases 2 and 3) emerge from a chemical increase in *δE* as emerge from a mechanical increase in *δE* (a rapid increase in length, ΔL, Fig. 4A). Figure 5C is a plot of phase 2 amplitudes obtained at different [P_i_]_f_ overlaid with experimental data (Dantzig et al. 1992). Figure 5D is a plot of phase 2 rates obtained at different [P_i_]_f_ overlaid with experimental data (Dantzig et al. 1992).

A least squares fitting routing was not used in the above analysis, and no attempt was made to refine fits to experimental data. For example, in all simulations changes in *a* were assumed to be single exponential (Eq. 9) even though more complex force-or time-dependences of *a* are justified and could be used to significantly improve fits to experimental data. Moreover, additional biochemical states in the actin-myosin ATPase reaction cycle exist that provide additional parameters that can be used to significantly improve fits to experimental data. The conclusion here is that while fits to time courses can be improved by expanding the model, the four thermodynamic phases of a force transient that emerge from a minimal binary mechanical model are qualitatively consistent with the four phases observed experimentally.

## DISCUSSION

Based on a minimal binary mechanical model (Fig. 1A) I have shown here that four well-defined thermodynamic phases of a muscle force transient emerge from a single two-state molecular mechanism. This is in contrast to popular corpuscular mechanic models in which a different molecular mechanism is required for each phase.

In 1938, based on careful measurements of muscle power and heat output, A.V. Hill developed a chemical thermodynamic model of muscle contraction. At the time little was known about molecular mechanisms of muscle contraction, but A.V. Hill knew that once discovered, his model provided the framework into which the “detailed machinery must be fitted” (Hill 1966).

However, in the 1950’s static electron micrographs of muscle revealed myosin crossbridges between actin and myosin filaments (Hanson and Huxley 1953; Huxley and Niedergerke 1954), inspring a corpuscular mechanic determinism that required a theoretical framework fundamentally different from chemical thermodynamics (Hill 1974). This corpuscularian framework provides the foundation for most models of muscle contraction to date.

According to Gibbs, thermodynamics is the laws of mechanics of systems of molecules as they appear to us, not the laws of mechanics of the molecules within a system as corpuscularians reason them to be (“rational mechanics”). To describe the phenomenally complex mechanics of molecules subject to thermal fluctuations within a system, we must on some scale consider average mechanical properties, and the only scale at which we can define a mechanical average is the scale on which mechanics are physically constrained and measured. Huxley assumed that myosin crossbridges were physically constrained by a crystalline lattice within muscle (Huxley 1957), but we now know that every component of this “lattice” thermally fluctuates along with every other molecule in muscle as one large macromolecular assembly.

In a chemical thermodynamic model, the laws of mechanics of muscle are macroscopically constrained and described by system free energy changes. With any reversible chemical step there is an infinitude of possible molecular mechanisms (thermal fluctuations of complex structures, different combinations of molecular steps, and rapid binding and detachment of ions, water and accessory proteins). Yet at a given standard condition the energetics and kinetics that describe all possible mechanisms by which an ensemble takes a chemical step are defined by a small number of macroscopic parameters such as chemical states, system free energy, system entropy (concentration gradients), a system force, and the extent, *a*, to which that force approaches the equilibrium isotherm.

These macroscopic parameters are not determined from rational molecular mechanisms; they are constrained by and determined from system free energy equations. Of course, molecular mechanisms can be inferred, but only after the macroscopic state variables that satisfy the laws of mechanics of the system (Eqs. 3 and 4) have been determined. For example, from the value of *a* > 1 determined from the amplitude of phase 2 force generation, we infer that work is performed by titin with this phase.

The transient analysis herein focuses entirely on a single, two-state binding mechanism from which four thermodynamic processes emerge. Phase 1 occurs with a rapid change, *δE*, in the system energy. Phase 2 occurs when adiabatic molecular force generation (Eq. 3) reaches a pseudo-equilibrium isotherm. Phase 3 occurs with a chemical relaxation toward the equilibrium isotherm (Eq. 9). And phase 4 occurs with a return to the initial force along the equilibrium isotherm. The goal here was to further demonstrate, using nothing more than basic physical principles and measured parameters, that the most minimal version of a binary mechanical model is consistent with experimental data. The conclusion is that while fits to time courses could be improved by expanding the model, the four thermodynamic phases of a force transient that emerge from a single binding mechanism are qualitatively consistent with the four phases observed experimentally.

The binary mechanical model in Fig. 1A reconciles disparate models of molecular motor mechanochemistry (Baker 2022b). The basic thermodynamic model, consistent with A.V. Hill’s analysis (Hill 1938), accounts for steady state muscle mechanics and energetics (Baker and Thomas 2000). Equations 3 and 4 describe the innumerable thermodynamic pathways along which muscle generates force (Fig. 1C, bottom to top), power output (Fig. 1C, right to left), work loops that perform net work on the surroundings (Fig. 1C, counter clockwise loops), and work loops that generate periodic dissipative forces (Fig. 1C, clockwise loops) (Baker 2022a). Here, I have shown that the four phases of a muscle force transient each correspond to a different thermodynamic pathway. Broadly, thermodynamic lessons learned from the most efficient machines on earth have provided new insights into chemical energetics and kinetics leading to the first explicit solution to the Gibbs paradox (Baker 2023). Specific to biology, this model pulls molecular biology away from an obsolete corpuscularian worldview back toward Bernoulli, Carnot, Boltzmann, Gibbs, and A.V. Hill down a path along which classic thermodynamics is applied to biological systems.

## METHODS

### Thermodynamic Analysis

Equations 3 (black lines) and 4 (colored curves) are plotted in Fig. 1C. Equation 4 is plotted at different ΔG° values (1 kT increments), and Eq. 3 is plotted at different ΔG° values (increments of 5). Each line represents a reversible binding reaction, and many complex mechanical behaviors resembling muscle mechanics emerge from these binding pathways.

### Computer Simulations

Using master equations 7, 8 and 9, the rate constants defined by Eq. 6, and model parameters in Table I, MatLab (Mathworks, Natick, MA) is used to simulate muscle force transients.

## AUTHOR CONTRIBUTIONS

JEB conceptualized and developed the theory, wrote and executed the code, designed and made the figures, and wrote the manuscript.

## ACKNOWLEDGEMENTS

I thank JWG, AVH, Julie, and the students, colleagues, and mentors who have over many years inspired and guided this work. This work was funded by a grant from the National Institutes of Health 1R01HL090938-01.

